# Exploring the mismatch between the theory and application of photosynthetic quotients in aquatic ecosystems

**DOI:** 10.1101/2022.09.07.507017

**Authors:** Matt T. Trentman, Robert O. Hall, H. Maurice. Valett

**Author notes:** **Author contribution statement:** MTT and ROH came up with the research question and designed the study approach. MTT and ROH conducted data simulations, and MTT, ROH, and HMV collected data. MTT, ROH, and HMV refined ideas and wrote the paper. **Data availability statement:** Data and metadata will be made available in the Dryad repository. The code for data simulations, analyses, and figure creation are available at https://github.com/mtrentma.

## Abstract

Estimates of primary productivity in aquatic ecosystems are commonly based on variation in O_2_, rather than CO_2_. The photosynthetic quotient (PQ) is used to convert primary production estimates from units of O_2_ to C. However, there is a mismatch between the theory and application of the PQ. Aquatic ecologists use PQ=1-1.4. Meanwhile, PQ estimates from the literature support PQ=0.1-4.2. Here, we describe the theory on why PQ may vary in aquatic ecosystems. We synthesize the current understanding of how processes such as NO_3_^−^ assimilation and photorespiration can affect the PQ. We test these ideas with a case study of the Clark Fork River, Montana, where theory predicts that PQ could vary in space and time due to variation in environmental conditions. Finally, we highlight research needs to improve our understanding of the PQ. We suggest departing from fixed PQ values and instead use literature-based sensitivity analyses to infer C dynamics from primary production estimated using O_2_.

**Scientific Significance Statement:** Accurate measures of primary production in aquatic ecosystems are necessary to quantify energy availability to higher trophic levels and biological effects on global CO_2_ concentrations, among other reasons. However, we commonly measure primary production using O_2_ because it is easier, despite our motivation to measure the rate of fixed C, and then use the photosynthetic quotient (the ratio of O_2_ release to CO_2_ fixed, PQ) to convert O_2_ based metabolism to CO_2_. This study provides a summary of the current mismatch between our current knowledge and the application of PQ, highlights our current knowledge gaps, and emphasizes the need to use literature-based sensitivity analysis rather than uninformed fixed PQ values.

## Introduction

Primary production in aquatic ecosystems fixes dissolved carbon dioxide (CO_2_) and subsequently releases dissolved oxygen (O_2_) via photosynthesis. Primary production can control the carbon (C) cycling in aquatic ecosystems, which has implications for the availability of energy to higher trophic levels, dissolved nutrient availability and cycling, and global climate dynamics due to the effects on atmospheric CO_2_ concentrations (McKinley et al., 2017). Primary production can be measured either via the rate of gaseous product (O_2_) generation or as the rate of reactant (CO_2_) consumption following the generic photosynthesis reaction (eq. 1) or by measuring the C assimilated into biota using isotopic approaches measured in microcosms (Nielsen, 1951; Peterson, 1999) to whole ecosystems (Bower et al., 1987).

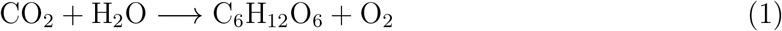

Most studies measure primary production as the change in O_2_, rather than CO_2_ assimilated in aquatic ecosystems, because O_2_ is easier to continuously and accurately measure using sensors. Indeed, much of the early research on aquatic ecosystem metabolic processes is based on models measuring the change of O_2_ with either diel variation (Odum, 1956) or experimental manipulation of light (Nielsen, 1951). Furthermore, methods to measure CO_2_ lagged that of O_2_, and also require knowing pH or alkalinity along with more difficult computation to account for the overall dissolved inorganic carbon (DIC) pool (Stets et al., 2017).

Because we are often more interested in C cycling we convert primary production measured with O_2_ data to C using the photosynthetic quotient (PQ), i.e., the molar ratio of the flux of O_2_ released to CO_2_ assimilated during photosynthesis (eqs. 2,3).

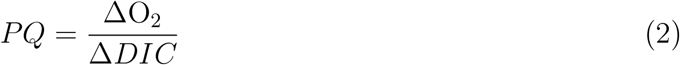

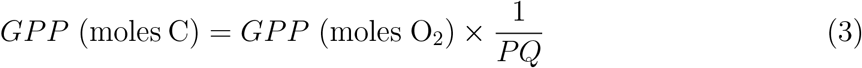

Despite its evident importance to translating from one chemical species to another, the PQ term has been dogmatically employed without appreciating the existing degree of uncertainty and its implications. A mismatch between theory and application of the PQ exists within and across aquatic ecosystem sub-fields. For example, it is common to assume a PQ of 1 or 1.2 for freshwater ecosystems, even though these values are not necessarily backed by measurements or theory. Marine researchers commonly use a value of 1.4, which is in part based on theory, but under specific assumptions (Williams et al., 1979). Fixed and field-specific PQ values are used despite PQ measurements from the literature ranging from 0.1 to 4.2 (Table 1).

**Table 1:** Measured PQ using varying methods and ecosystems show a broad range of possible PQ. SI Table 1 contains a detailed citation and summary table.

| **Aquatic Ecosystem** | **Method** | **Number of Studies** | **Range of PQ** |
| --- | --- | --- | --- |
| Marine | $^{14}\text{C}$ | 6 | 0.1-4.2 |
|  | DIC | 4 | 0.4-1.6 |
| Freshwater | $^{14}\text{C}$ | 2 | 0.3-2.0 |
|  | DIC | 3 | 0.13-1.4 |

Physiological mechanisms that co-occur with oxygenic carbon fixation alter the rate of net O_2_ production because they occur at rates and scales commensurate with that employed by our methods even as those methods fail to distinguish them. For example, photorespiration and nitrate (NO_3_^−^) assimilation are net sinks and sources of O_2_ (respectively) that are rarely accounted for in models that estimate primary production (Figure 1). These oversights may have extensive and unappreciated consequences for our understanding of C fluxes and rates of energy flow associated with primary production across ecosystems.

**Figure 1:**
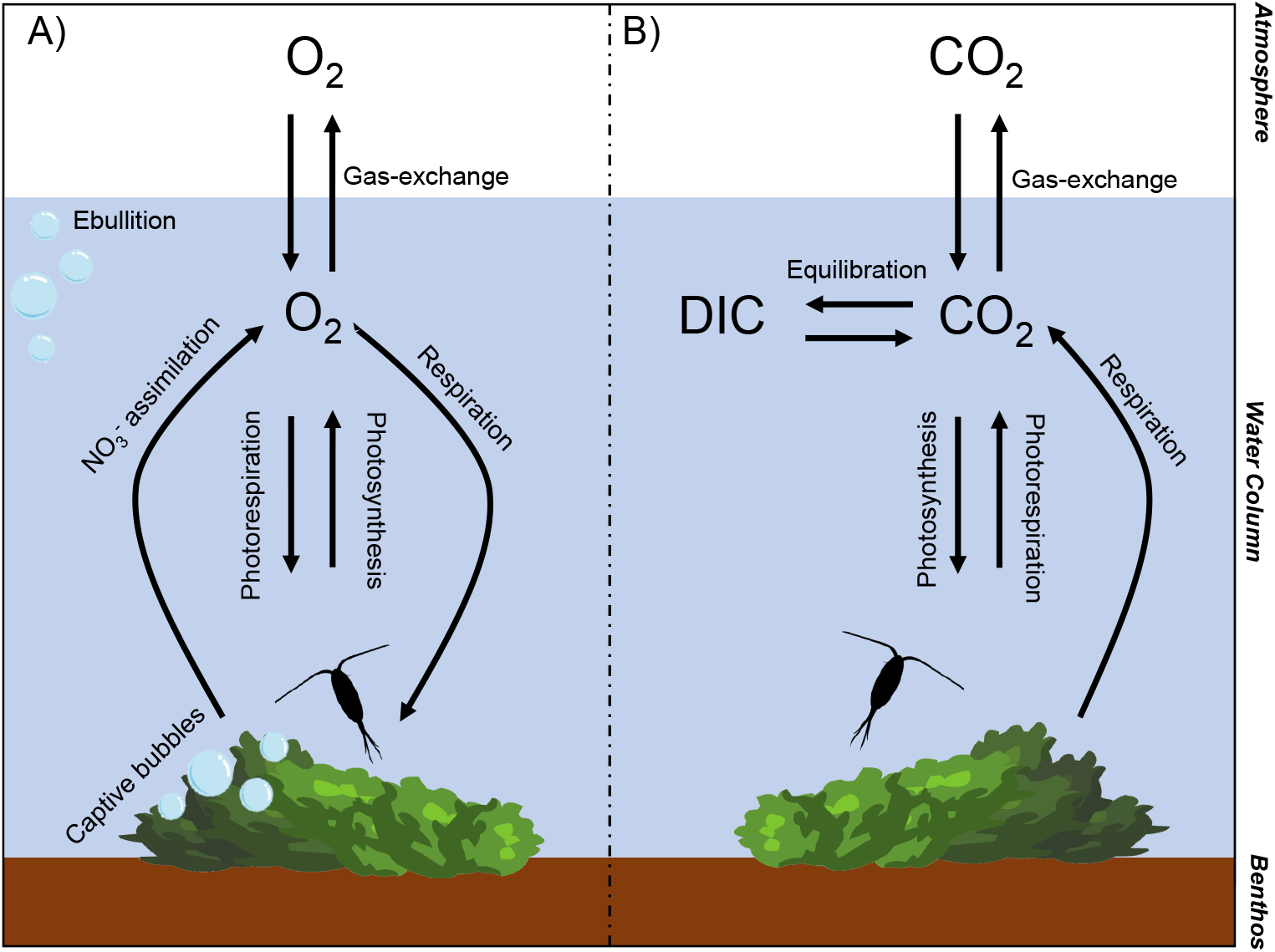
Summary of processes relative to the primary production of benthic algae and phytoplankton that affect dissolved O_2_ (A) and CO_2_ (B). Both O_2_ and CO_2_ are affected by photosynthesis and respiration, while O_2_ is uniquely affected by NO_3_^−^ assimilation and photorespiration. O_2_ concentrations are much more likely to be affected by captive bubbles or ebullition since O_2_ has low solubility above saturation. In contrast, CO_2_ equilibration with dissolved inorganic carbon (DIC) reduces the potential for bubble formation. Note that the magnitude of the effect of any given process on dissolved gas concentrations depends on a variety of environmental conditions specific to that process.

Here, we explore the current status of PQ values in aquatic ecosystems and their historical application, with the goal to enable better estimates of C transformations based on O_2_ production. Specifically, we will:

1. Summarize measurements of PQ within and between marine and freshwater aquatic ecosystems,
2. Describe the state of chemical and biological theory addressing how and why PQ varies across species and environmental conditions,
3. Provide a case study of a river where theory predicts that PQ should vary in space and time based on environmental conditions and algal succession, and
4. Highlight research needs to improve our understanding of how we translate between C and O metabolism.

Overall, we suggest departing from minimally informed fixed PQ values and instead use literature-based sensitivity analyses to inform how PQ choice affects interpreting C dynamics from primary production estimated using O_2_. Furthermore, we encourage simultaneous measurements of O_2_ and CO_2_ when possible, to provide empirical evidence of environmental PQ variation and accurate elemental translation.

We recognize from the outset that PQ is difficult to assess because we cannot directly measure photosynthesis. Measurements of photosynthesis will always co-occur with respiration and parsing the two measurements is problematic; it is common to measure respiration (in the dark), then measure net ecosystem production (in the light) and subtract respiration from net ecosystem production. Thus, variation in respiration, and in particular, respiration quotients (RQ) will affect our assessment of PQ. In this work, we are not considering the effects of RQ, since a summary of the RQ variation with environmental conditions already exists (Berggren et al., 2012). Even so, RQ and PQ should both be considered when interpreting O_2_ and CO_2_ metabolism data.

### History of PQ measurements and values used across aquatic ecosystems

Most of our knowledge of the PQ comes from experiments and measurements in marine ecosystems on planktonic algae. While our purpose is not to exhaustively review the literature, the papers we encountered were mostly marine with few from freshwater, including only one study with measurements of PQ on freshwater benthic algae (Table 1). The bias of PQ measurements towards phytoplankton and marine ecosystems may be due to historic method constraints on the accurate measurement of CO_2_ (Williams and Robertson, 1991). Instead, radioactive ^14^C has been used since at least the 1960s (Strickland, 1960) to measure primary productivity (referred to here as ^14^C productivity). Given the complexities of using radioactive materials in the natural environment, this method is much more favorable at small scales (e.g., bottles that capture microscopic primary producers) and in the laboratory, thus the initial bias towards phytoplankton and marine ecosystems. However, the ^14^C approach is not directly comparable to measures of O_2_ evolution because the incubation results in something between gross primary production and net primary production (Peterson, 1980). These uncertainties in the measurement of C fixation bias estimates of the PQ.

Williams and Robertson (1991) review showed PQ ranged between 0.5-3.5, with most measurements made based on ^14^C productivity. They attributed the broad range in PQ to the ^14^C productivity problem (Peterson, 1980), and suggested that a more realistic range of values for phytoplankton is between 1.1-1.4 (Williams and Robertson, 1991). This range was confirmed by Laws (1991), who used chemical equations of photosynthetic products, and reported estimates of phytoplankton chemical composition to show that PQ=1.1 to 1.4. These values of PQ represent broad averages, and more contemporary site-specific studies using more accurate methods to measure primary production (i.e., direct measurement of CO_2_ or dissolved inorganic carbon (DIC) compared to ^14^C productivity) have resulted in PQ values that differ from 1.1-1.4 based on specific species and environmental conditions (Table 1). Despite the known variation of the PQ, using a fixed value of 1.35-1.45 is still common in marine ecology (Bolden et al., 2019).

Freshwater ecologists also commonly use fixed PQs but usually employ values of 1 or 1.2, based more on tradition rather than on theory or data. We know of no direct source suggesting the use of PQ=1 although it is probably safe to assume that this value reflects only the photosynthetic creation of glucose (C_6_H_12_O_6_), which results in molar equivalence between O_2_ released and CO_2_ fixed. The assumption of glucose as the only photosynthetic product oversimplifies organic matter synthesis and ignores other processes that can affect dissolved O_2_ concentrations. The PQ value of 1.2 similarly does not have a common source, although some attribute this value to Wetzel (2001) or Bott et al. (1978), despite neither source providing convincing evidence for a fixed value. Although the range of reported PQ measurements in fresh waters is between 0.13 and 2.0 (Table 1), we have not found an article that applies a fixed value other than 1 or 1.2 when the PQ isn’t directly measured. Moreover, none have accounted for the implications associated with the uncertainty of the PQ values employed.

We suggest that aquatic ecologists need to use greater scrutiny when applying PQ values to translate O_2_-based metabolism to rates of C fixation. Given the need for an accurate PQ, we summarize knowledge of why the PQ may vary across species and environmental conditions, and use this knowledge to generate hypotheses about the biological and environmental conditions causing the PQ to vary. While most of this knowledge is based on studies of phytoplankton from marine ecosystems, the same concepts often hold in freshwater ecosystems.

### PQ theory

PQ can be separated into multiple components, with each dictated by the characteristics of the species of primary producer and the environmental conditions surrounding the organism (Williams and Robertson, 1991). The appropriate PQ value that represents biological effects on O_2_ and CO_2_ (*PQ*_*Bio*_) can be represented by eq. 4:

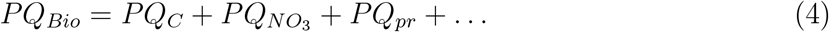

where *PQ*_*C*_ refers to the change in O_2_ and CO_2_ associated with photosynthesis, 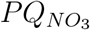 refers to the release of O_2_ associated with the reduction of nitrate (NO_3_^−^) for nitrogen (N) assimilation, and *PQ*_*pr*_ is the reduction of O_2_ associated with photorespiration. The ellipsis (…) represents the myriad of other biological processes that may affect O_2_ or CO_2_ availability (e.g., nitrification, sulfate reduction). We focus on the three described components identified by considerable evidence to measurably affect the PQ in most aquatic ecosystems.

#### PQ_C_

The *PQ*_*C*_ represents the change in O_2_ and CO_2_ associated with photosynthesis. The *PQ*_*C*_ can be broken down into the various products of photosynthesis dictated by the state of reduction of the products for the general compound *C*(*H*)_*Y*_ (*O*)_*Z*_ (eq. 5) (Ryther, 1956; Laws, 1991):

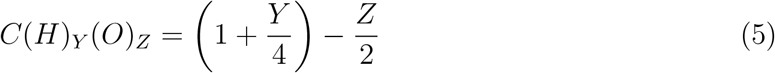

where *Y* and *Z* represent the number of H and O atoms in the photosynthetic product, respectively. Thus, the net change in O_2_ and CO_2_ associated with the sum of *C*(*H*)_*Y*_ (*O*)_*Z*_ products equals *PQ*_*C*_. The most common products intimately coupled with photosynthesis are glucose (*C*_6_*H*_12_*O*_6_), glycolic acid (*C*_2_*H*_3_*O*_3_), saturated fatty acids (e.g., *C*_20_*H*_30_*O*_2_), nucleic acids (e.g., *C*_38_*H*_47_*O*_28_*N*_15_*P*_4_), and protein (e.g., *C*_18_*H*_24_*O*_6_*N*_5_). The proportion of C in each of these compounds relative to the total C in the algal cell will dictate the value of the *PQ*_*C*_. The range of plausible 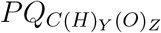 values is 0.625-1.5, with the lowest value assuming the production of glycolic acid only, and the highest value assuming the sole production of saturated fatty acids. In reality, these extremes are not likely, and the *PQ*_*C*_ will fall somewhere in the middle of this range. Laws (1991) estimated the ‘typical’ planktonic cell to contain 40% each of proteins and carbohydrates, 15% lipids, and 5% nucleic acids, which equates to a *PQ*_*C*_ of 1.08. This value is specific to phytoplankton, and there is likely species-specific variation in *PQ*_*C*_ based on algal physiology and life history. Notably, benthic algae may have a different *PQ*_*C*_ than phytoplankton given the need for making structures to anchor to the substratum (among other physiological differences); however, there is little information available to calculate the *PQ*_*C*_ of benthic algae.

Few papers have measured the effect that specific photosynthetic end products have on the PQ, although speculation abounds that environmentally driven changes in lipid production increase the PQ. These changes in the PQ are likely associated with variation in light and energy availability occurring in environments replete with other possible limiting nutrients as the algal cell uses the opportunity to generate lipids that require more energy and C, thus increasing the PQ (Palmucci et al., 2011; Norici et al., 2011). Indeed, many studies have observed increased PQ with diel increases in light (Carvalho, 2014), and experimental light manipulation (Du et al., 2018). The mechanistic link (or lack thereof) between environmental variables and lipid production is a knowledge gap relevant to estimates of PQ in freshwater and marine ecosystems.

#### 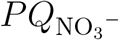

The biosynthesis associated with primary production also requires N, and the concurrent reduction of NO_3_^−^ by the cell for assimilation will release O_2_, thus increasing the PQ. Specifically, 2 moles of O_2_ are released for every mole of NO_3_^−^ reduced to NH_3_.

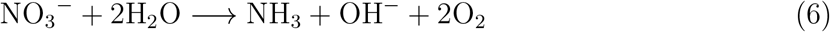

Organismal demand for N controls 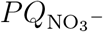; C:N ratios in organismal biomass are a good indicator of this demand with lower values reflecting greater mass-specific demand for N. Finally, it is energetically favorable for organisms to assimilate N as ammonium (NH_4_^+^), which does not produce O_2_. Therefore, the relative amount of NO_3_^−^ in the total dissolved inorganic nitrogen pool (NO_3_^−^ + NH_4_^+^; DIN) also controls the amount of O_2_ released as a part of N assimilation (Smith et al., 2012).

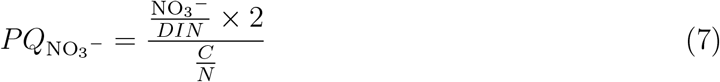

Lab experiments demonstrate that O_2_ release due to NO_3_^−^ assimilation increases PQ values (Raine, 1983; Bell and Kuparinen, 1984). Raine (1983) manipulated NO_3_^−^ concentrations of salt water containing phytoplankton in the lab, while keeping all other environmental conditions constant. PQ varied from 1.0-2.25 and the measured change in PQ was similar to the theoretical change in PQ based on chemical stoichiometry (e.g., eq. 7)(Raine, 1983). Field measurements tell the same story; Smith et al. (2012) observed a 0.18 increase in the PQ over ∼ 30-y in a Rhode Island estuary. The observed increase in the PQ coincided with increased proportion of DIN present as NO_3_^−^ associated with a new wastewater treatment plant upstream of the estuary. The change in relative N availability (i.e., eq. 7) suggests that the PQ should have increased by 0.15-0.23, which includes the observed increase of 0.18 (Smith et al., 2012). The match between expected values derived from chemical stoichiometry and empirical measurements enhances confidence in estimates of 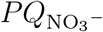 compared to *PQ*_*C*_. The lowest confidence is assigned to estimates of *PQ*_*pr*_, which addresses the complicated and poorly addressed influences of photorespiration.

#### PQ_pr_

Photorespiration is the light-dependent O_2_ reduction and CO_2_ release by photoautotrophs. Unlike mitochondrial respiration, photorespiration only occurs in light, does not conserve energy in ATP, and does not use substrates from the tricarboxylic acid cycle (Tolbert, 1974). Instead, photorespiration reduces O_2_ via the RuBisCO mediated reaction with Ribulose 1,5-bisphosphate (RuBp) that generates C-2 compounds (e.g., 2-phosphoglycolate) further processing of which consumes more O_2_ and liberates CO_2_.

While the end result of this process includes the release of CO_2_, some or all of this CO_2_ is ‘recycled’, or immediately fixed by photosynthesis, rather than being released from the cell (Søndergaard and Wetzel, 1980). Photorespiration was initially referred to as a ‘wasteful process’ given that glycolate is not needed and is often excreted when in excess (Tolbert and Zill, 1956). Later studies have discovered that glycolate is eventually funneled through multiple reactions to contribute to the Calvin-Benson cycle, although at a higher energy cost than via photosynthesis (Moroney et al., 2013). During photorespiration O_2_ occupies the RuBisCO binding site, limiting CO_2_ fixation and reducing rates of net photosynthesis. Photosynthesis inhibition at high O_2_ concentrations is a common phenomenon explained by photorespiration. Most algal species have developed C concentrating mechanisms to build up DIC around the RubisCO enzyme, limiting photorespiration (Giordano et al., 2005). Aquatic photoautotrophs need C concentrating mechanisms because they commonly face an environment with low CO_2_ concentrations, and need to build up CO_2_ near RubisCO for photosynthesis (Moroney et al., 2013). Despite the evolution of C concentrating mechanisms to limit its effects, photorespiration is still necessary. For example, mutant cyanobacteria missing some photorespiration pathways could not to survive outside of abnormally high CO_2_ environments (Eisenhut et al., 2008), evidence that photorespiration is necessary despite C concentrating mechanisms.

While the purpose and potential benefits of photorespiration are not yet fully understood (Osmond and Grace, 1995; Shi and Bloom, 2021), generally 3 moles of O_2_ are reduced for every mole of CO_2_ produced during photorespiration, which ultimately lowers PQ (Stewart, 1974). Multiple experiments have shown that photorespiration constitutes 20-40% of net photosynthesis across a variety of aquatic ecosystems and measurement scales (Burris, 1981; Parkhill and Gulliver, 1998; Kliphuis et al., 2011; Buapet et al., 2013), particularly when O_2_ is readily available and CO_2_ is relatively low (Vance and Spalding, 2005). Photorespiration rates are often ∼ 0 below 100% O_2_ saturation, low at or below 100% O_2_ saturation, and highest when O_2_ is super-saturated. Although, low CO_2_ (and indirectly elevated pH) can often increase photorespiration rates as long as some O_2_ is present (Kliphuis et al., 2011). Multiple studies have measured PQ *<* 1, which is often attributed to photorespiration (Hough, 1974; Burris, 1981; Pokorny et al., 1989). Furthermore, some studies have experimentally manipulated light (Iriarte, 1999) or O_2_ concentrations (Burris, 1981) and found PQ *<* 1 matching the theoretical effect of photorespiration. We note that photorespiration has not been observed (when it otherwise should have been) under some environmental conditions and with some specific algal species (Lloyd et al., 1977; Birmingham et al., 1982).

### Towards predicting when and where PQ may vary based on environmental conditions

PQ estimates from the literature and direct experimentation provide a theoretical basis for hypothesizing when and where PQ may vary with environmental conditions. Here, we synthesize the relevant information to aid researchers in addressing variation in the PQ. Literature supports the assertion that 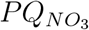 and *PQ*_*pr*_ are likely to have the strongest effect on the PQ, and these effects can be predicted based on the proportion of NO_3_^−^ that is DIN, the C:N ratio of the algal biomass, and the O_2_ % saturation. We used data from the literature to simulate the effect of environmental conditions on the PQ using eq. 7 (for effects of 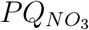) and data from Burris (1981) (for effects of the *PQ*_*pr*_) (Figure 2). We conducted the simulations across the plausible range of each variable in the environment, where the environmental conditions ranged for the proportion of DIN that is NO_3_ from 0 and 1, for O_2_ saturation from 40 and 180%, and with an algal C:N composition of 5 or 20. This simulation provides a broad hypothetical framework for predicting when and where the PQ may vary. Results from the simulation (Figure 2) illustrate the potential for *PQ*_*pr*_ to increase the *PQ*_*Bio*_ by as much as 0.8 when dissolved O_2_ is low, and decrease by as much as 1.2 when O_2_ is well above saturation. Changes associated with 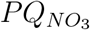 are less dramatic, with changes in the *PQ*_*Bio*_ of 0.2 when N demand is high (low biomass C:N, Figure 2a) and less when N demand is low (high biomass C:N, Figure 2B). We note that the simulated changes in *PQ*_*Bio*_ are relative to an assumed *PQ*_*C*_ of 1.1, an assumption that carries its own degree of uncertainty. While the literature supports the possibility of variation in the *PQ*_*C*_, we do not believe there is enough information to adequately predict anything other than a fixed *PQ*_*C*_.

**Figure 2:**
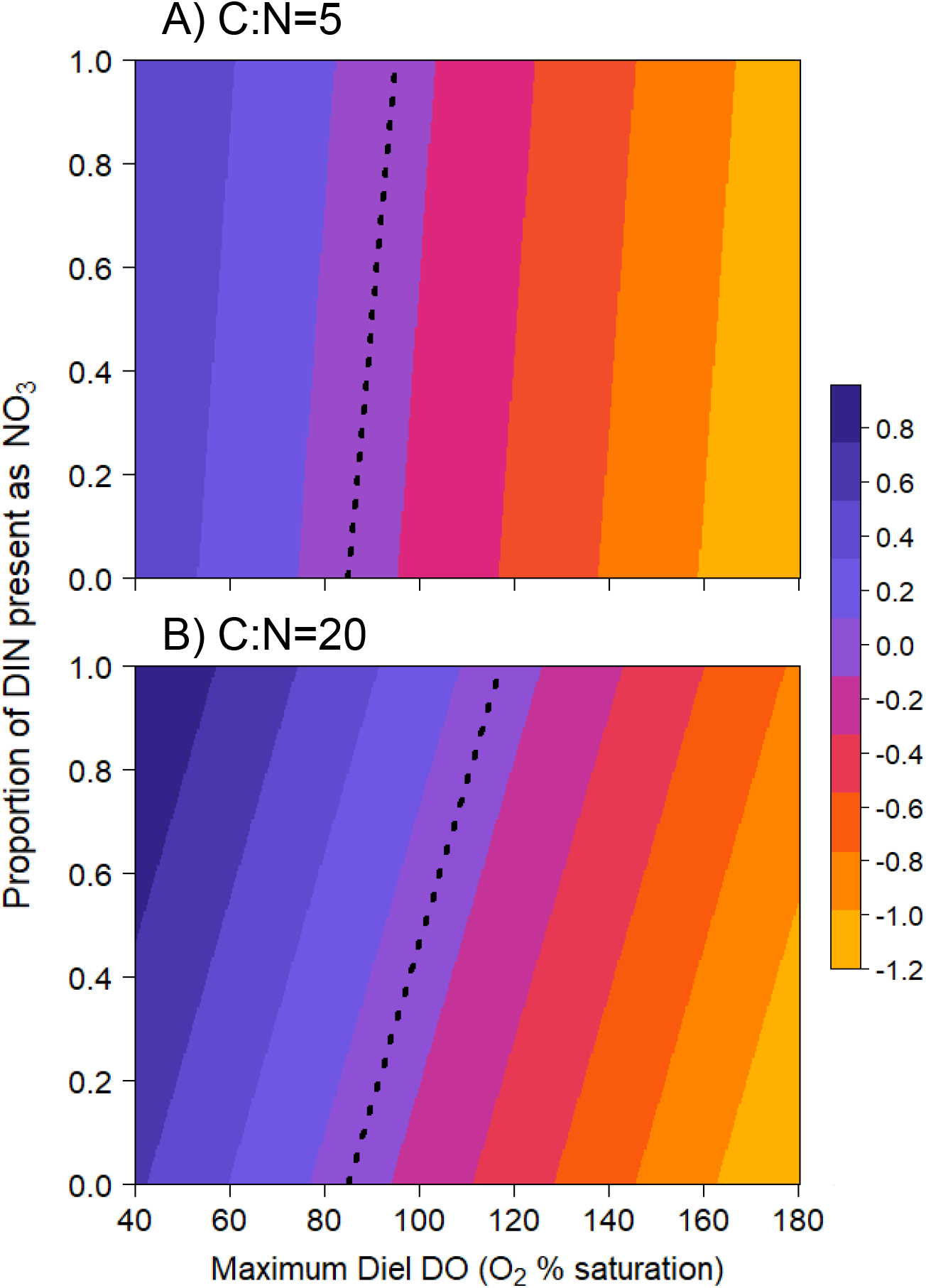
The simulated change in PQ associated with the 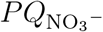 (y axis) and the *PQ*_*pr*_ (x-axis) components. The top panel assumes a fixed C:N of 5, while the bottom panel assumes a fixed C:N of 20. Both simulations assume *PQ*_*C*_=1.1 and the dashed line shows the combination of environmental conditions where PQ does not vary from 1.1. The PQ will increase (positive change relative to *PQ*_*C*_) when NO_3_^−^ is a high proportion of DIN and the aquatic ecosystem is below 100% O_2_ saturation. In contrast, the PQ will decline the most when NO_3_^−^ is a low proportion of DIN and waters are supersaturated with O_2_. Finally, when the demand for N is low (indicated by high C:N ratio), the effect of 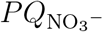 on the PQ change is less than when N demand is high.

Primary productivity varies with light intensity, and thus PQ can change with the diel progression of light (Hough, 1974; Iriarte, 1999; Mercado et al., 2003). Furthermore, the degree to which the PQ will vary during a day will depend on the rate of primary productivity, given that ecosystems with low productivity are unlikely to supersaturate with O_2_, limiting the potential effect *PQ*_*pr*_ may have on *PQ*_*Bio*_. The proportion of NO_3_^−^ that is DIN can vary seasonally or spatially, particularly in lotic ecosystems. Accordingly, we simulated how different fixed rates of gross primary production (GPP), in units of moles O_2_ released (*GPP*_*O*_), interact with environmental conditions to control the *PQ*_*Bio*_.

Specifically, we simulated three scenarios: 1) low GPP rates (i.e., O_2_ saturation never exceeds 100%) and low NO_3_^−^ assimilation rates (assuming mineralization of NH_4_^+^ as is likely at low productivity rates), 2) high GPP rates (i.e., O_2_ saturation exceeds 100% during most of the photic period) and low NO_3_^−^ assimilation rates (assuming a low proportion of NO_3_^−^ that is DIN), and 3) high GPP rates and high NO_3_^−^ assimilation rates (assuming a high proportion of NO_3_^−^ that is DIN). We again assume that *PQ*_*C*_ is fixed at 1.1. We simulated the diel O_2_ saturation curves assuming low and high fixed values for GPP (19 and 250 mmol m^*−*3^ d^*−*1^, respectively) and ecosystem respiration scaled with GPP, while gas-exchange, average depth, temperature, and pressure were held constant (SI 1, SI Table 2). We used the theory presented above to predict 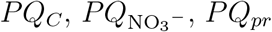, and *PQ*_*Bio*_ in each scenario. We then used the *PQ*_*Bio*_ derived from each scenario to calculate GPP, in units of fixed C (*GPP*_*C*_). Finally, we compared the *GPP*_*C*_ predicted with the *PQ*_*Bio*_ from each scenario with the *GPP*_*C*_ estimated using fixed PQ values commonly used in marine and freshwater ecosystems, and from the range observed in the literature.

In low productivity ecosystems, N assimilation and photorespiration are likely low, thus the *PQ*_*Bio*_ ≈ *PQ*_*C*_ (Figure 3). Based on our current understanding of *PQ*_*C*_, when *PQ*_*Bio*_ ≈ *PQ*_*C*_ the choice of traditional fixed PQ values will minimally bias estimates of *GPP*_*C*_ from measured *GPP*_*O*_. In contrast, highly productive ecosystems with low NO_3_^−^ will have a *PQ*_*Bio*_ that is *<* 1. This scenario causes the largest bias of *GPP*_*C*_ estimates from *GPP*_*O*_. PQ values *<* 1 will have a higher bias than values *>* 1 of the same magnitude (i.e., 0.8 compared to 1.2) (Figure SI 1). In the final scenario, the O_2_ consumed through photorespiration is negated by the addition of O_2_ from NO_3_^−^ assimilation, resulting in a *PQ*_*Bio*_ *> PQ*_*C*_. In this scenario, the *PQ*_*Bio*_ is similar in magnitude with the traditional fixed values, with minimal bias based on the choice of *PQ*_*Bio*_.

**Figure 3:**
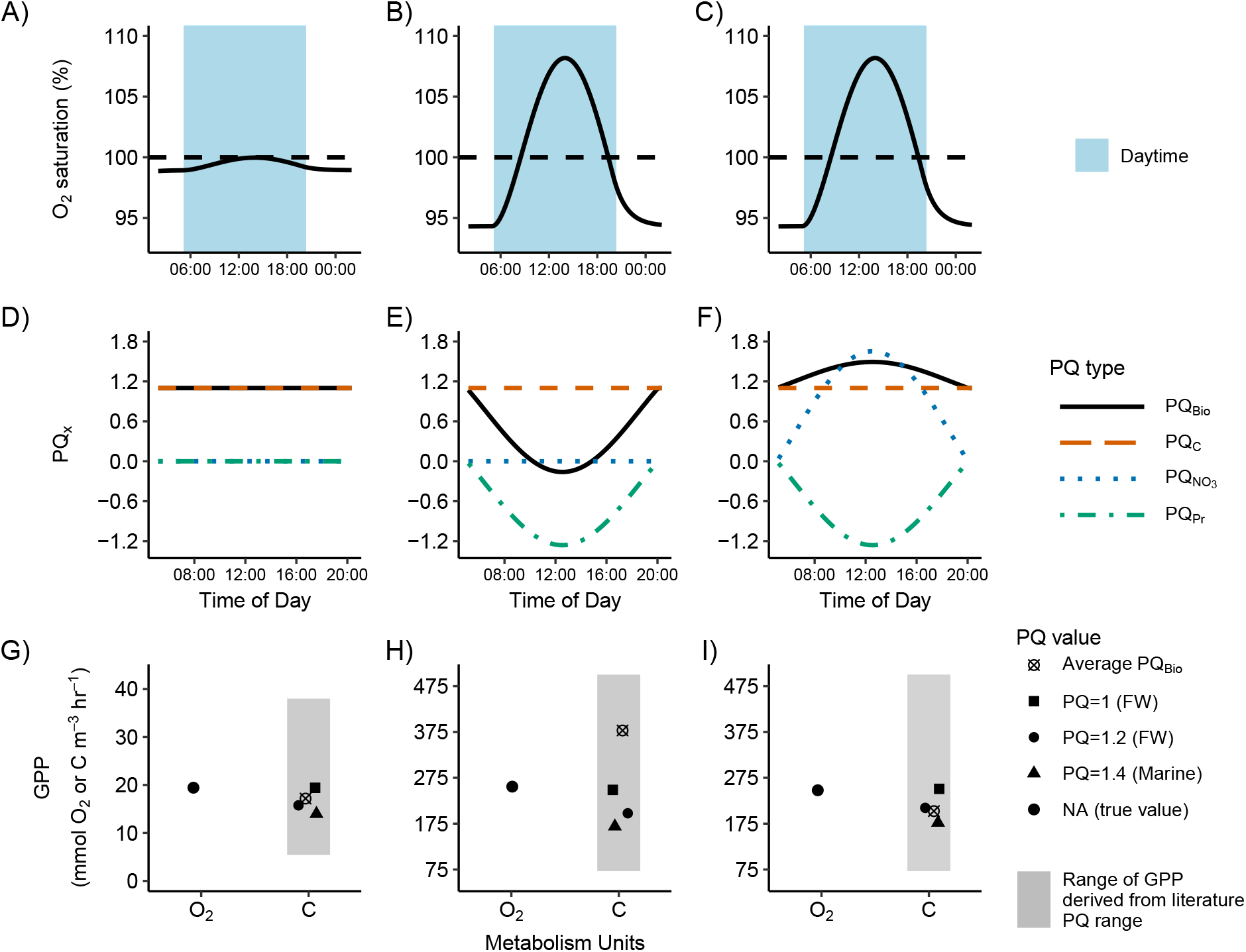
The simulated effect of each *PQ*_*x*_ component on the *PQ*_*Bio*_ given different *GPP*_*O*_ rates and NO_3_^−^ availability. Each column represents a different scenario: left column-low GPP and low NO_3_^−^, middle column-high GPP and low NO_3_^−^, right column-high GPP and high NO_3_^−^. The O_2_ saturation data are simulated based on fixed values of GPP (A=19 mmol O_2_ m^*−*3^ hr^*−*1^ and B,C= 250 mmol O_2_ m^*−*3^ hr^*−*1^). SI 1 contains full methods on O_2_ saturation simulation. The blue shaded area represents the daylight period. The middle row shows the simulated effects of each *PQ*_*x*_ component on the *PQ*_*Bio*_ (Eq. 4). The mean *PQ*_*x*_ for each scenario was fixed based on our current knowledge of the effect of environmental conditions on each compartment. In the low GPP scenario (D), *PQ*_*Bio*_ ≈ *PQ*_*C*_ since the effects of *PQ*_*Pr*_ and 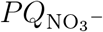 are negligible when O_2_ saturation is *<* 100% and NO_3_ concentration is low. For the high productivity and low NO_3_^−^ scenario (E), the *PQ*_*Bio*_ is *< PQ*_*C*_ due to photorespiration occurring when O_2_ saturation is ≥ 100%. For the high productivity and high NO_3_^−^ scenario (F), *PQ*_*Bio*_ *> PQ*_*C*_ primarily controlled by the large effect of 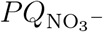. The final row shows how the fixed GPP value in units of O_2_ would be transformed to C units using: the mean *PQ*_*Bio*_ from D-F, the two most common fixed freshwater (FW) values (1,1.2), the most common marine value (1.4), and a conservative range of PQ values observed in the literature (0.5, and 3.5, respectively). The error associated with using the conventional fixed PQ values is low when GPP rates are low, and vice versa when rates are high. The highest error when converting GPP from units of O_2_ to C occurs when GPP rates are high, NO_3_^−^ is low, and photorespiration is occurring.

### Considerations of observation error and measurement bias

The above theory provides testable hypotheses that can be confronted with measurements of the PQ (*PQ*_*Meas*_) across environmental conditions. In an ideal world, *PQ*_*Meas*_ would match *PQ*_*Bio*_; however, this scenario is unlikely ever true. It is necessary to consider the observation error (*PQ*_*Obs error*_) and measurement bias (*PQ*_*Meas bias*_).

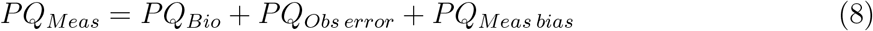

*PQ*_*Obs error*_ is the error imposed by imperfectly measuring O_2_ or DIC. For example, an improperly calibrated or drifting sensor may add observation error to the PQ. In general, it is possible to limit *PQ*_*Obs error*_ with careful calibration and lab work. Even so, there are statistical approaches to account for measurement error (e.g., state-space models, Pedersen et al. 2011; Appling et al. 2018). We define *PQ*_*Meas bias*_ as the error imposed when a model for dissolved O_2_ or DIC does not reflect the process of interest. *PQ*_*Meas bias*_ is process error and can occur for a variety of reasons. For example, abiotic processes other than the biological processes associated with primary production can bias estimates of GPP, (e.g., bubble formation in chambers). *PQ*_*Meas bias*_ will be most prevalent when comparing photosynthesis based on O_2_ or CO_2_ at the ecosystem level and provides a substantial challenge for discerning the biological signal within *PQ*_*Meas*_. While *PQ*_*Meas bias*_ can take many forms, we highlight a few sources of *PQ*_*Meas bias*_ that deserve further consideration and research.

For ecosystems with attached benthic primary producers (Figure 1A), captive bubbles (i.e., bubbles from primary production trapped in biofilms) are common, but understudied in productive ecosystems and are more likely to affect O_2_ than CO_2_. Dissolved CO_2_ readily equilibrates with the DIC pool in many circumneutral pH environments, while the CO_2_ remaining as dissolved gas has a much higher solubility than O_2_. Bolden et al. (2019) attribute observed captive bubbles on marine coral reefs to abnormally low PQ measurements after ruling out the contribution of *PQ*_*pr*_ and 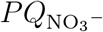. The bias in measured O_2_ was large enough that the authors suggested that O_2_ based estimates of metabolism are unreliable in highly productive benthic environments (Bolden et al., 2019). Ebullition occurs when captive bubbles are released into the water column, and eventually traverse the air-water divide. The effect of ebullition on primary production estimates has been documented in many lakes, where emissions of bubbles can be similar in magnitude to diffusive O_2_ exchange (Koschorreck et al., 2017), and contain up to 20% of daily net oxygen production (Howard et al., 2018). Ultimately, captive and ebullated bubbles can affect estimates of *PQ*_*Bio*_ in variable ways depending on the degree to which gases in bubbles equilibrate with the water around them (Hall Jr and Ulseth, 2020).

Groundwater inputs can affect O_2_ and CO_2_ concentrations in receiving freshwater ecosystems (Sear et al., 1999; Duvert et al., 2018). Generally groundwater in local flow is replete with CO_2_ and low in O_2_ due to soil respiration (Tank et al., 2018); low O_2_ groundwater can bias estimates of GPP (Hall Jr. and Tank, 2005). The contribution of groundwater CO_2_ can generally be predicted by stream hydrology, with higher CO_2_ contributions in small headwater streams compared to larger rivers (Hotchkiss et al., 2015; Horgby et al., 2019*b*); however, knowledge of local land use and geology, among other factors, are needed to adequately predict groundwater O_2_ and CO_2_ contributions (Tank et al., 2018; Horgby et al., 2019*a*; Hutchins et al., 2021). Furthermore, groundwater contributions are not static and will likely vary with environmental conditions (Ulseth et al., 2018). Overall, groundwater inputs of CO_2_ are likely to be high in many freshwater ecosystems, which may lead to an underestimation of *PQ*_*Bio*_.

### Case study: The Upper Clark Fork River, MT

The literature clearly supports that the *PQ*_*Bio*_ can vary from the canonical fixed values based on environmental conditions with measures further complicated by a mismatch between *PQ*_*Bio*_ and *PQ*_*Meas*_. Next, we provide a case study of a river where changing environmental conditions predict that the *PQ*_*Bio*_ varies in space and time, and differs from *PQ*_*Meas*_. This case study reveals the inability to reconcile the application of canonical PQs, PQ theory, and measurements of PQ.

The Upper Clark Fork River in western Montana is a mid-order river that is highly productive during the summer growing season due to ample light, warm water, and plenty of nutrients (both N and P). The NO_3_^−^ concentration is high near the headwaters of the river, due to a combination of natural and anthropogenic inputs, and generally declines moving downstream. Phosphorus (as soluble reactive phosphorus, SRP) is high in the middle section of the Clark Fork due to weathering of P-rich rocks in this region (Carey, 1991). Variability in nutrients and productivity make the Clark Fork a useful location to explore how environmental conditions may translate to variation of the *PQ*_*Bio*_ in space and time. Specifically, we explore how variation in NO_3_^−^, and the maximum daily O_2_ saturation might control spatial and temporal variation in the *PQ*_*Bio*_ during the summer growing season.

We present data collected in the Clark Fork from summer 2020 (1 Aug - 30 Oct) at two sites, an upstream site near Warm Springs, Montana (Site 1) and a downstream site near Clinton, Montana (Site 2). Data presented are: O_2_ concentrations collected using continuous sensors (PME) at 10-min. intervals, and DIN concentrations (as NO_3_^−^ + NH_4_) collected as bi-weekly filtered grab samples and analyzed using standard methods. The O_2_ data were used to calculate maximum O_2_ saturation for each day, and estimate *GPP*_*O*_ using a single-station approach. More details on data collection and analyses are provided in SI 3. We use these data and the theory presented above to predict *PQ*_*Bio*_ at each site over the algal growing season. Data presented here are unpublished and associated with long-term efforts to characterize the relationship between nutrients, algal biomass, and metabolism in the Clark Fork (H.M. Valett, unpublished data).

Diel O_2_ reflected a highly productive river with large daily excursions. Maximum daily O_2_ saturation exceeded 100% at both sites for most of the time series (Figure 4). Maximum daily O_2_ saturation dipped below 100% at the downstream site for the final 2 wk, and at the upstream site for the final 3 d of data. Our current understanding of the link between O_2_ saturation above 100% and photorespiration suggests that photorespiration was likely affecting *PQ*_*Bio*_. The effect of photorespiration likely declined in the later portion of the sampling period as max O_2_ saturation dipped below 100%, with a longer dip at the downstream site. The proportion of DIN that was NO_3_^−^ was generally higher (between 0.6-0.95) at the upstream site, compared to the downstream site (between 0.2-0.9) (Figure 4). At both sites, the proportion declined by 0.4 during the growing season and then rebounded towards the end of the growing season. The predicted effect of O_2_ release from NO_3_^−^ assimilation on the PQ was greater at the upstream site compared to the downstream site, with similar amounts of within-site variation over the growing season (Figure 4).

**Figure 4:**
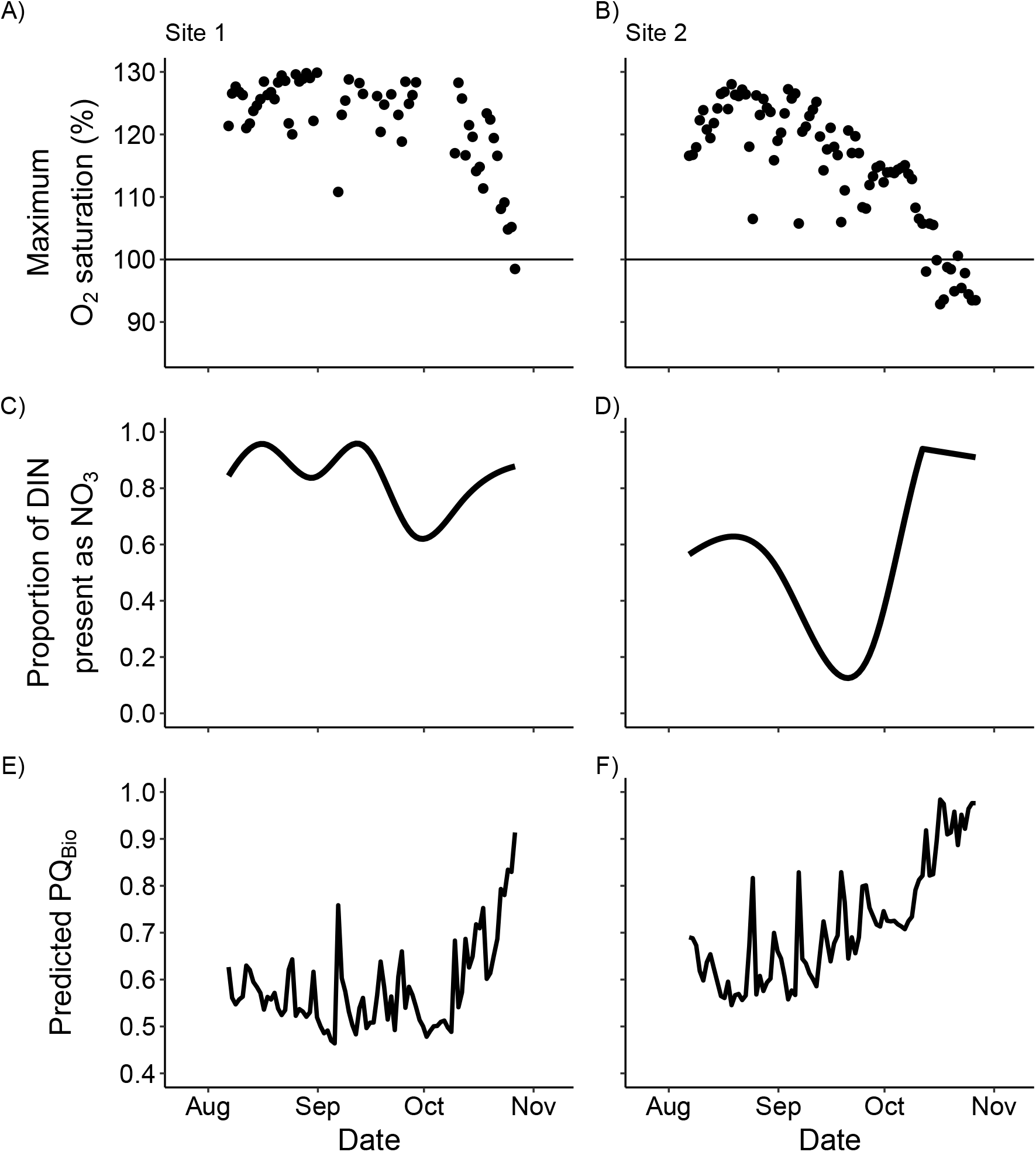
The measured environmental conditions that may affect *PQ*_*Bio*_ from two sites on the Upper Clark Fork River, MT during summer 2020: The maximum O_2_ saturation from continuous (10 min) diel measurements (A,B), and the proportion of DIN present as NO_3_^−^ from bi-weekly linearly interpolated grab samples (C,D). Measured environmental conditions predicted *PQ*_*Bio*_ *<* 1 through time at each site (E,F).

We applied the predicted *PQ*_*Bio*_ reflecting the influences of the components addressed herein to the *GPP*_*O*_ measurements derived at the upstream and downstream sites to estimate *GPP*_*C*_ (Figure 5). We then compared *GPP*_*C*_ rates estimated with the predicted *PQ*_*Bio*_ to the *GPP*_*C*_ estimated with the canonical PQ value of 1.2, as well as a conservative range of PQ values from the literature (0.4-3.5). At both sites, *GPP*_*O*_ rates declined by 50-60% during the measurement period. During the early part of the measurement period, *GPP*_*C*_ using the predicted *PQ*_*Bio*_ was ∼ 30% greater than the *GPP*_*C*_ estimated using the traditional fixed PQ of 1.2. The difference between estimates declined as the growing season progressed. At Site 1, the two estimates converged for the final 2 weeks of measurements as max O_2_ saturation declined to less than 100% saturation and *PQ*_*Bio*_ increased to near 1. At site 2, the two estimates of *GPP*_*C*_ nearly converged for the final few days of measurements, again matching patterns in maximum O_2_ saturation. Assuming our predicted *PQ*_*Bio*_ is accurate, using a fixed value of 1.2 may lead to an error in *GPP*_*C*_ estimates of 0-33% based on the site and time of the year.

**Figure 5:**
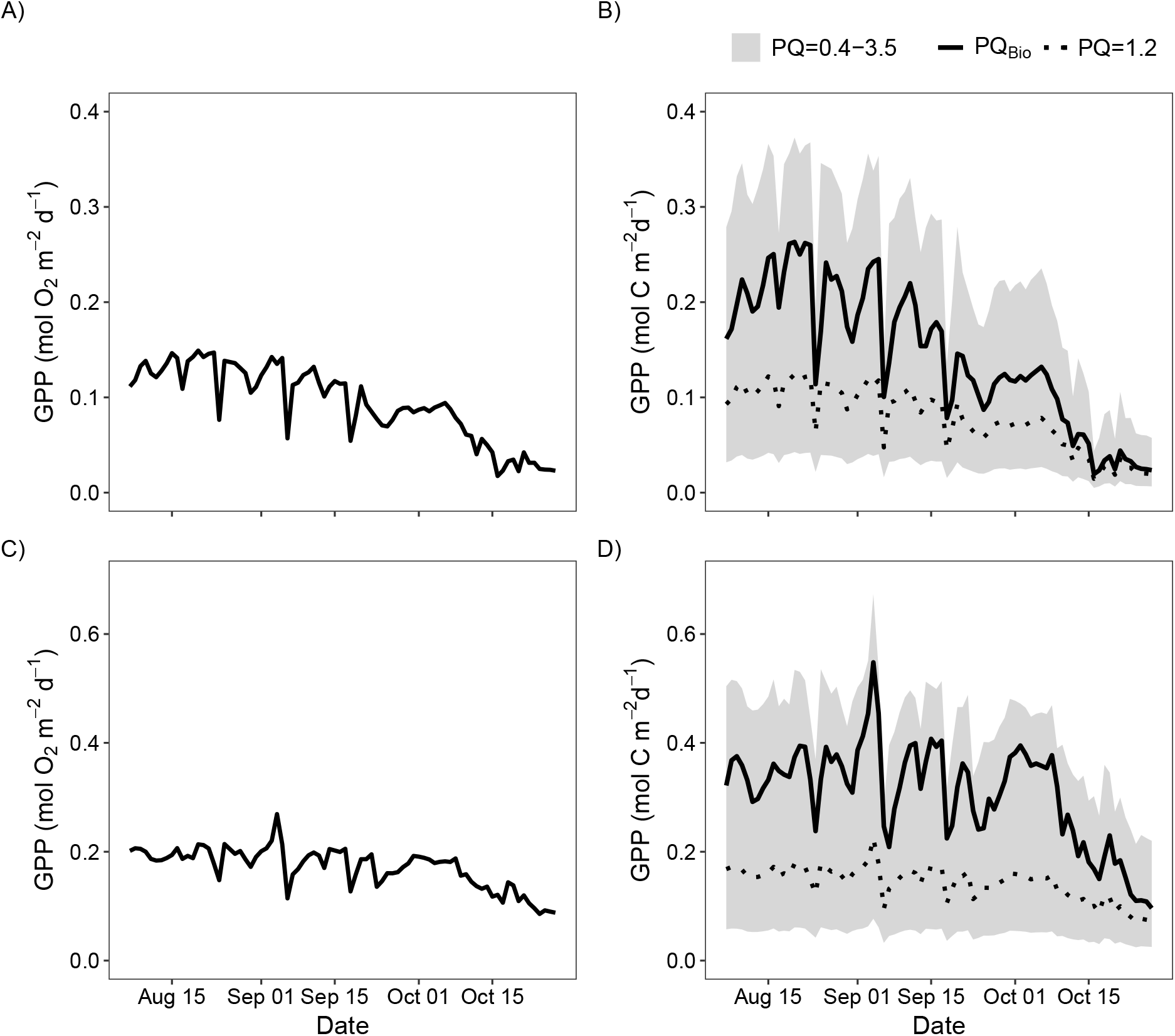
The measured *GPP*_*O*_ (in units of O_2_; A,C) and estimated *GPP*_*C*_ (in units of C); B,D) from two sites in the UCFR in the summer of 2020 (Site 1, top; Site 2, bottom). We used multiple PQs to estimate the *GPP*_*C*_, including the *PQ*_*Bio*_ from Figure 4, PQ=1.2 (traditionally used in freshwater ecosystems), and the range of PQ values from the literature (0.4-3.5). The *GPP*_*C*_ estimates using *PQ*_*Bio*_*are*0-33% higher than *GPP*_*C*_ estimates based on the fixed value of 1.2.

To capture the potential uncertainty imposed by the various PQs used here, we also compared the *GPP*_*C*_ estimated from the canonical value with the range of PQs found in the literature. The *GPP*_*C*_ using the range of literature PQ values was ± 63% different than the *GPP*_*C*_ estimates using PQ=1.2 (Figure 5), which is a wide range that has implications for how we interpret and analyze *GPP*_*C*_ estimates.

Finally, to employ a more controlled assessment of *PQ*_*Bio*_, we compared the *PQ*_*Bio*_ predicted from environmental conditions with a set of measured PQ values using instruments to directly quantify CO_2_ and O_2_. We used sealed recirculating acrylic chambers (Dodds and Brock, 1998) to measure the PQ from substrate collected from the Clark Fork River in summer 2021. The PQ was estimated using the change in O_2_ (measured from continuous sensors) and DIC (estimated from CO_2_ measured using *in situ* sensors (DeGrandpre, 1993) and *ex situ* measurements of alkalinity). Six PQ measurements were made based on independent substrata and at varying levels of productivity, provided by altering the available light using a shade cloth. SI 3 contains a detailed description of the methods. Based on linear regression of molar rates among the six experiments, the *PQ*_*Meas*_ from the sealed chambers 1.6 (± 0.5 st. dev., Figure 6).

**Figure 6:**
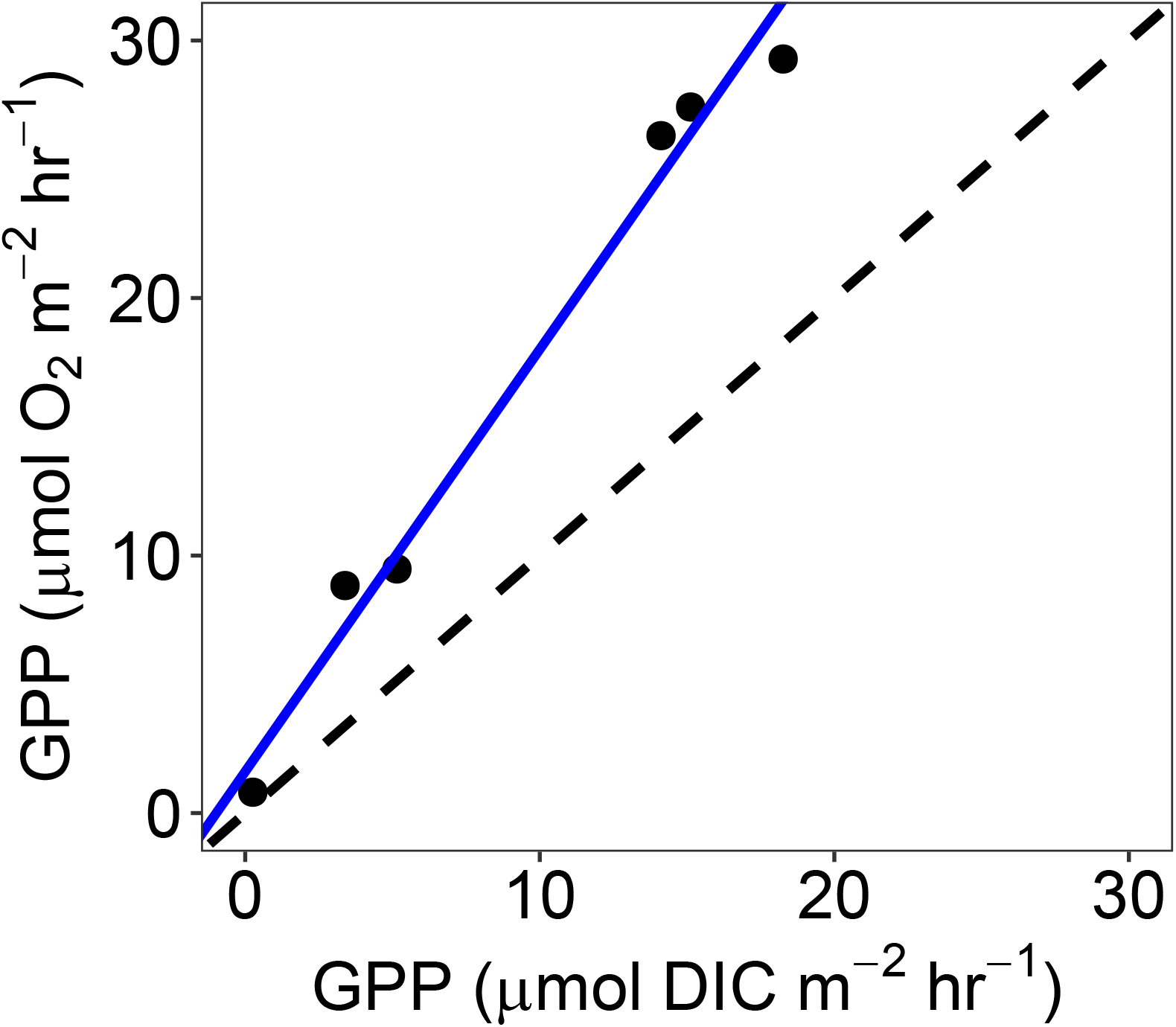
The relationship between O_2_ and DIC-based gross primary production measured simultaneously in a chamber using substrate from the Clark Fork River. The slope of the blue line was calculated using a reduced major axis approach since the axes are symmetrical and represents *PQ*_*Meas*_=1.6 (± 0.5 st. dev.), which is higher than the predicted *PQ*_*Bio*_ from the same river.

The difference between the mean predicted *PQ*_*Bio*_ (0.7) and the mean *PQ*_*Meas*_ (1.6) in the Clark Fork River shows that there were either substantial bias and error in our measures of C and O_2_ metabolism (i.e., *PQ*_*Meas*_) or poor predictions based on gaps in the current theory (i.e., *PQ*_*Bio*_). These data cannot adjudicate the true value of the PQ and rather exposes our extensive knowledge gaps in theory and measurement of PQ. It is necessary to resolve these problems in PQ theory and measurements in order to generate accurate assessments of *GPP*_*C*_.

### Knowledge gaps

#### Photorespiration

Photorespiration may be more prevalent when estimating aquatic primary production than we know. Studies that estimate photorespiration at the ecosystem level are uncommon (but see Parkhill and Gulliver (1998)), despite ample evidence that photorespiration is occurring in highly productive and pH-neutral waters. Thus, more ecosystem-level estimates of the prevalence and magnitude of photorespiration are necessary. While some evidence shows a relationship between photorespiration and O_2_ saturation or the ratio of O_2_ to CO_2_, the data representing these empirical relationships is lacking. Furthermore, it is not clear if empirical relationships are species-specific. For example, some species of red and brown algae suppress photorespiration more than many green algae species (Reiskind et al., 1989). In general, it is not clear how strongly species differ in the magnitude and prevalence of photorespiration, and if the potential variation is relevant to ecosystem-level estimates of primary production.

#### NO_3_^−^ assimilation

While multiple field and lab studies have characterized the effect of O_2_ release during NO_3_^−^ assimilation on the PQ, our ability to predict this effect is more complicated than our current knowledge (i.e., Eq. 7). Eq. 7 does not account for the scenario where NH_4_^+^ and NO_3_^−^ are both saturated in respect to nutrient demand (e.g., algae are limited by their maximum nutrient uptake rate rather than nutrient availability, Dodds et al. (2002)). For example, a realistic scenario in many agricultural streams is that both NH_4_^+^ and NO_3_^−^ are present in high concentrations (e.g., 10 *μ*M and 110 *μ*M, respectively; M.T. Trentman, personal observation). In this scenario, using Eq. 7 would lead to an overestimate of the effect of NO_3_^−^ assimilation on the PQ because the form of N uptake is dominated by NH_4_^+^, which is preferentially assimilated by organisms. In short, Eq. 7 should account for the abundance of NH_4_^+^ in relation to demand as well as the relative availability of NO_3_.

#### Photosynthetic products

Information on the production of different C products during photosynthesis are uncommon, specifically for freshwater algae and species that have no commercial interest, making it difficult to calculate the theoretical 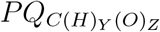. Thus, our understanding of how *PQ*_*C*_ may vary by species and with environmental conditions (e.g., light) is limited. However, most photosynthetic products, including proteins, nucleic acids, and glucose, have a 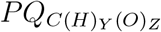 of approximately 1. That leaves glycolic acid and saturated fatty acids as two possible sources of variation, which could decrease and increase the 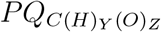, respectively, based on their content in an algal cell. It is unlikely that the production of glycolic acid is substantial enough to shift the PQ given the relatively low production rate (Cheng et al., 1972). It is more likely that the species-specific variability in saturated fatty acid production (Harwood, 2019) would change the 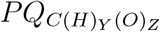. For example, The lipid content of benthic algae can range in complexity (i.e., O and H content, Guschina and Harwood (2006); Li-Beisson et al. (2019)), but there are little data on how this variation in lipid chemistry affects the proportion of the total C of algal biomass. Research is needed that quantifies the rate of saturated fatty acid production across algae species, and how this production varies with environmental variability.

#### More measurements of PQ, particularly on freshwater benthic algae

More measurements of PQ, particularly on freshwater benthic algae and using methods other than ^14^C, would improve our understanding of how the *PQ*_*Bio*_ varies among species and with environmental variables. The PQ can be measured through a variety of approaches that can be generally separated by the location of the measurements, whether controlled (e.g., in bottles or chambers) or in the natural environment. Measuring the PQ requires simultaneous monitoring of the C that is fixed and the O_2_ that is released during the process of primary production (Although, see the above text that describes the measurement challenges imposed by respiration). The measurement of O_2_ can be accurately obtained with low-cost optical sensors (assuming no captive bubbles). Estimating the change in DIC is more involved, and includes either direct measurements of DIC or is accomplished by estimating the DIC pool via measures of dissolved CO_2_ concentrations and either pH or alkalinity (Dickson et al., 2007). In highly buffered waters the relative decline in DIC will be small, and this decline is difficult to accurately quantify, but the decline in CO_2_ will be large. Given this fact, we briefly highlight the pros, cons, and limitations of the best-known approaches that use accurate measures of O_2_ and CO_2_.

Using sealed containers to measure algal productivity allows for control of environmental conditions. Controlled measurements of PQ will promote equivalence between *PQ*_*Meas*_ and *PQ*_*Bio*_, since it will be easier to limit *PQ*_*Meas bias*_. The bottle approach is useful for phytoplankton, while more complex chambers should be used for benthic algae, especially from rivers. Unidirectional flow in rivers is one of many controls on primary productivity, thus chambers with continuous, circulating flow are needed to measure the PQ for riverine benthic algae (e.g, Dodds and Brock (1998); Rüegg et al. (2015). PQ measurements sealed from the atmosphere are best because, if used properly, the chambers allow the researcher to rule out gas-exchange between the water and the atmosphere, which can be challenging to measure in low productivity and turbulent ecosystems (among other conditions, Hall Jr and Ulseth (2020)). The benefits of bottles or chambers should be balanced with the potential bias of measuring *ex situ* processes. When working with chambers, it is common to not let O_2_ exceed 100% saturation in order to limit bubble formation in the chamber. Thus, chamber experiments may be biased in that the effects of *PQ*_*Pr*_ are limited. Finally, large chambers can be expensive and cumbersome to use (M.T. Trentman pers. obs.).

Measuring diel variation in O_2_ and DIC to estimate metabolic processes is another approach to estimate the PQ, although in most scenarios *PQ*_*Meas*_ ≠ *PQ*_*Bio*_ due to *PQ*_*Meas bias*_. The bias of PQ measured with diel sensors will be particularly problematic given that there is a range of environmental controls on reach scale or whole lake O_2_ and DIC budgets. Bubbles, groundwater inputs, and the complexities of DIC chemistry (e.g., calcite precipitation), among many other factors, may impose higher variation on *PQ*_*Meas*_ than the biological signal. That said, simultaneous monitoring of diel O_2_ and CO_2_ concentrations is a relatively new approach made possible by the increased availability of accurate CO_2_ sensors (Vachon et al., 2020). However, at the time of writing this article, the models used to estimate metabolism from DIC are still in development despite their prior use (Wright and Mills, 1967). A degree of caution is necessary to interpret reach-level estimates of PQ given that measurements will likely not reflect *PQ*_*Bio*_. Future methods and models should be developed with the specific goal of parsing *PQ*_*Bio*_ from *PQ*_*Meas*_ at all levels of measurement.

## Conclusions

An accurate *PQ*_*Bio*_ is needed for understanding C metabolism derived from O_2_ data. The current evidence suggests that the *PQ*_*Bio*_ can vary with environmental conditions. Thus, the use of fixed canonical *PQ*_*Bio*_ values is likely causing inaccurate estimates of C-based metabolism from O_2_ data. While the current theory provides a basic understanding of the processes that may control the *PQ*_*Bio*_, much research will be necessary to use environmental conditions to predict *PQ*_*Bio*_ values more accurately and precisely. Furthermore, consideration needs to be given to observation error and measurement bias that will be common when measuring the PQ in the natural environment. Until we improve PQ theory and our ability to reconcile error and bias from *PQ*_*Meas*_, we suggest we should employ more scrutiny when applying PQ values to convert O_2_ based metabolism to C. Specifically, researchers should move beyond fixed values of the PQ. Instead, we suggest matching our current knowledge of how the PQ varies with the environmental conditions at any given measurement site to generate a range of plausible PQ values (e.g., Figure 5) from the literature. Thus, accounting for the uncertainty of the PQ when estimating C-based metabolism from O_2_ data.

## Acknowledgements

National Science Foundation EPSCoR Cooperative Agreement OIA-175735, National Science Foundation grants EF-1834679 and LTREB DEB 1655197 supported our research. Amanda Spencer and Sam Bosio assisted with chamber PQ measurements.

## Notes

https://github.com/mtrentma

### Competing Interest Statement

The authors have declared no competing interest.

