## Supplemental Material for "Exploring the mismatch between the theory and application of photosynthetic quotients in aquatic ecosystems"

SI Table 1: Summary of studies that have measured the PQ cited in this paper. We did not conduct an exhaustive literature search; however, these papers likely represent the bulk of PQ research. The range of measured PQ are separated by study, aquatic ecosystem, and the method employed, where $^{14}$C represents primary production measured using $^{14}$C isotopes, and DIC represents primary production estimates using direct measurements of DIC or indirect estimates of DIC from a combination of $CO_2$, pH, and alkalinity.

| **Aquatic Ecosystem** | **Citation** | **PQ** | **Estimation approach** |
|---|---|---|---|
| *Marine* | | | |
| | Laws (1991) | 1.1, 1.4 | Measured |
| | Burris (1981) | 0.1-1.8 | Measured ($^{14}C$) |
| | Williams and Robertson (1991) | 1.1, 1.34 | Theoretical |
| | Smith et al. (2012) | 1.24, 1.42 | Measured ($^{14}C$) |
| | Du et al. (2018) | 0.9-1.4 | Measured (DIC via pH and TA) |
| | Bolden et al. (2019) | 0.47-1.02 | Measured (DIC and TA) |
| | Raine (1983) | 1.0-2.25 | Measured ($^{14}C$) |
| | Williams et al. (1979) | 1.1-2.25 | Measured ($^{14}C$) |
| | Mercado et al. (2003) | 0.9-1.6 | Measured (DIC via pH and TA) |
| | Iriarte (1999) | 0.7-4.2 | Measured (DIC and $^{14}C$) |
| | Rosenberg et al. (1995) | 0.4-1.0 | Measured (DIC via pH and TA) |
| | Carvalho (2014) | 0.13-2.55 | Measured ($^{13}C$) |
| *Freshwater* | | | |
| | Bell and Kuparinen (1984) | 1.63 | Measured ($^{14}C$) |
| | Pokorny et al. (1989) | 0.13-1.14 | Measured (DIC via pH and TA) |
| | Hanson et al. (2003) | 1.25 | Measured ($CO_2$) |
| | Sakamoto et al. (1984) | 0.9-2.0 | Measured (DIC and $^{14}C$) |
| | Kliphuis et al. (2011) | 0.3-1.4 | Measured (DIC via pH and TA) |
| | Davies and Hecky (2005) | 1.1 | ??? |

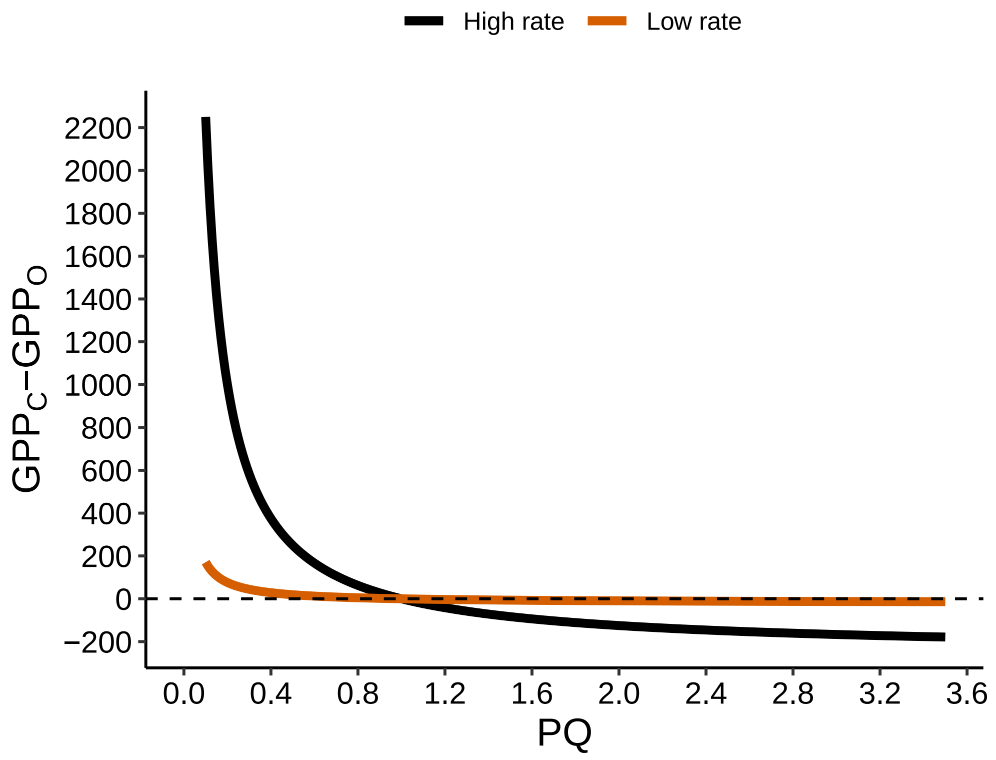


SI Figure 1: Simulated data showing how the difference between measured $GPP_O$ and estimated $GPP_C$ varies with the PQ. The data were simulated assuming a fixed $GPP_O$ value of 19 or 250 mmol $m^{-3}$ $d^{-1}$, and the range of PQ values represents the range of measured PQ values from the literature. PQ values $< 1$ will have a stronger effect on $GPP_C$ estimates than PQ values $> 1$ of the same relative magnitude from 1 (e.g., 0.8 vs 1.2).

### SI 1: Simulating $O_2$ data from known metabolic parameters

We simulated three scenarios that are representative of how diel variation in environmental conditions might affect the PQ.

- Scenario 1: Low GPP where $O_2$ saturation never exceeds 100% and low $NO_3^-$ assimilation.
- Scenario 2: High GPP where $O_2$ saturation exceeds 100% during most of the photic period and low $NO_3^-$ assimilation assuming a low proportion of $NO_3^-$ that is DIN.
- Scenario 3: High GPP where $O_2$ saturation exceeds 100% during most of the photic period and high $NO_3^-$ assimilation assuming a high proportion of $NO_3^-$ that is DIN.

For each scenario, we simulated a 26 hr $O_2$ time series at 10 minute intervals by choosing the metabolic parameters of gross primary production (GPP), ecosystem respiration (ER), and gas-exchange (K) that were realistic (i.e., found in the literature) and represented the desired conditions set for each scenario. In short, we chose metabolic parameters, and back-calculated the raw $O_2$ data that would generate the known parameters using the model of Hall et al. (2016). Light, average reach depth, temperature, and pressure are also needed to simulate the $O_2$ time series. Light at each time point was modeled using equations of (Yard et al., 2005) and the latitude, longitude, and standardized longitude of the Flathead Lake Biological Station, MT (47, 114, and 105, respectively). We held constant all other abiotic variables to avoid the influence of abiotic processes on the simulated $O_2$ data (SI Table 1). Thus, the simulated diel $O_2$ data only represent the effect of light and biological processes.

SI Table 2: Parameters used for each scenario to simulate $O_2$ data from known metabolic parameters. Note that GPP, ER, and K are fixed parameters, while barometric pressure, average reach depth, and temperature represent the values provided at each time point.

| **Parameter** | **Scenario 1** | **Scenario 2** | **Scenario 3** |
|---|---|---|---|
| GPP (mmol $O_2$ $m^{-3}$ $hr^{-1}$) | 19 | 250 | 250 |
| ER (mmol $O_2$ $m^{-3}$ $hr^{-1}$) | -47 | -250 | -250 |
| K ($d^{-1}$) | 15 | 15 | 15 |
| barometric pressure (mmHg) | 760 | 760 | 760 |
| average reach depth (m) | 1 | 1 | 1 |
| temperature (°C) | 15 | 15 | 15 |

### SI 2: Methods for collecting and modeling GPP estimates from the Upper Clark Fork River

We collected $O_2$ and temperature data at two points in the Upper Clark Fork River, MT. The first site (Site 1) is located near Warm Springs, MT (46.208494, -112.767485), and the second site (Site 2) is located near Clinton, MT (46.721546, -113.572918). We deployed PME dissolved oxygen sensors from Aug 1-Oct 30 2020 and measured $O_2$ and temperature at 10 minute intervals. We checked the sensors for $O_2$ concentration accuracy (using measurements at 100% saturation) before and after the deployment. There was minimal drift during the deployment. The average depth of the reach was fixed throughout the time period based on three average depth measurements during the deployment (one measurement in each month). While using a fixed depth is not typically ideal, there was relatively consistent discharge during the sampling period (Site 1 nearest USGS gage: 12323800, and Site 2 nearest USGS gage: 12331800), and consistent depths. The mean depth of the three measurements ($\pm$ standard deviation) was $0.60 \pm 0.04$ m for Site 1 and $0.64 \pm 0.05$ m for Site 2. Light was simulated based on (Yard et al., 2005). Elevation corrected barometric pressure was used from the Missoula International Airport (NOAA station id: 727730-24153).

We used these data to model daily estimates of gross primary production (GPP), ecosystem respiration (ER), and gas-exchange (K). We used the single-station open-channel (Odum, 1957) approach within the multi-day and hierarchical Bayesian model 'streamMetabolizer' (Appling et al., 2018) using the general equation:

$$\frac{dO_{2,t}}{dt} = GPP_t + ER_t + K_t \times D \tag{1}$$

where $\frac{dO_{2,t}}{dt}$ is the rate of change of $O_2$ at time point $t$, and $D$ is the deficit between actual and quilibrium concentrations of $O_2$ (Hall et al., 2016; Appling et al., 2018). We used the streamMetabolizer R package with the model variant 'b_Kb_oipi_tr_plrckm.stan' (Appling et al., 2018) and default priors in streamMetabolizer, which are normal or lognormal distributions created with the average and standard deviation of GPP, ER, and K from a meta-analysis compiled by (Hall et al., 2016). We solved for the model parameters using four Markov Chain Monte Carlo (MCMC) chains run on four cores with 500 burn-in and 500 saved steps.

We evaluated the model fit by checking that the chains converged (i.e., $\widehat{R} < 1.01$), and excluded estimates where chains did not converge. We evaluated the potential for equifinality for estimates of K and ER (Appling et al., 2018) by evaluating the relationship between K and ER. The slope of this relationship was not different from zero, suggesting no presence of equifinality. Finally, we removed estimates with incorrect signs (e.g., positive ER estimates).

### SI 3: Methods and data for measuring and analyzing the PQ in chambers

We measured the PQ in portable experimental chambers similar to that of Dodds and Brock (1998). The chamber consists of two main pieces, a main acrylic box for stream substrates that is transparent to light, and a PVC arm and propeller system that circulates water through the acrylic box, mimicking stream flow (Figure 2). We modified the chamber to pump water from the chamber via a submersible pump to make in-line measurements of $O_2$ (via a YSI dissolved oxygen probe attached to a sample box) and $CO_2$ (via a Submersible Autonomous Moored Instrument-$CO_2$ sensor (DeGrandpre, 1993) connected to the tubing with the manufacturers attachment). We used the chamber to measure the PQ on substrate from the Clark Fork River in downtown Missoula in August of 2021. On each day, the substrate were transported to the Flathead Lake Biological Station. A total of 21 PQ measurements were collected; however, due to sensor malfunction, only 5 measurements passed QAQC tests. For each chamber run, we placed substrate in the chamber, then we filled the chamber with water collected from the Clark Fork River that day. After sealing the chamber, we made sure that all air bubbles were removed from the in-line dissolved gas sample loop and from the chamber. The chamber was then placed in a 1100L cattle tank filled with water from Flathead Lake to reduce temperature fluctuations. We monitored the temperature of the tank water and used a submersible pump to replace water as needed to keep a constant temperature. We then initiated measurements of $O_2$, $CO_2$, and water temperature first in the dark (to measure ER), then in the light (to measure Net Ecosystem Productivity (NEP). We conducted three consecutive measurements on each substrate per day. We used the $CO_2$ data and the average alkalinity from periodic measurements (alkalinity did not change during the incubation) to estimate the dissolved inorganic C (DIC) pool (Dickson et al., 2007; Koschorreck et al., 2021). We then calculated the slope of the line of either $O_2$ or DIC during the ER and NEP measurements. We then used the following equation to calculate the oxygen-based GPP ($GPP_O$) and carbon-based GPP ($GPP_C$) using eq. 2

$$GPP_x \, slope = NEP_x \, slope + ER_x \, slope \tag{2}$$

where $x$ refers to either measurements of $O_2$ or estimates of DIC. Note that $ER_x$ slope is negative in eq. 2. Finally, we calculated PQ for each chamber run using the slope of six measurements of $GPP_x$ (eq. 3). We used a reduced major axis regression to estimate the slope given the axes are symmetrical.

$$PQ = slope \; of \frac{GPP_O}{GPP_{DIC}} \tag{3}$$

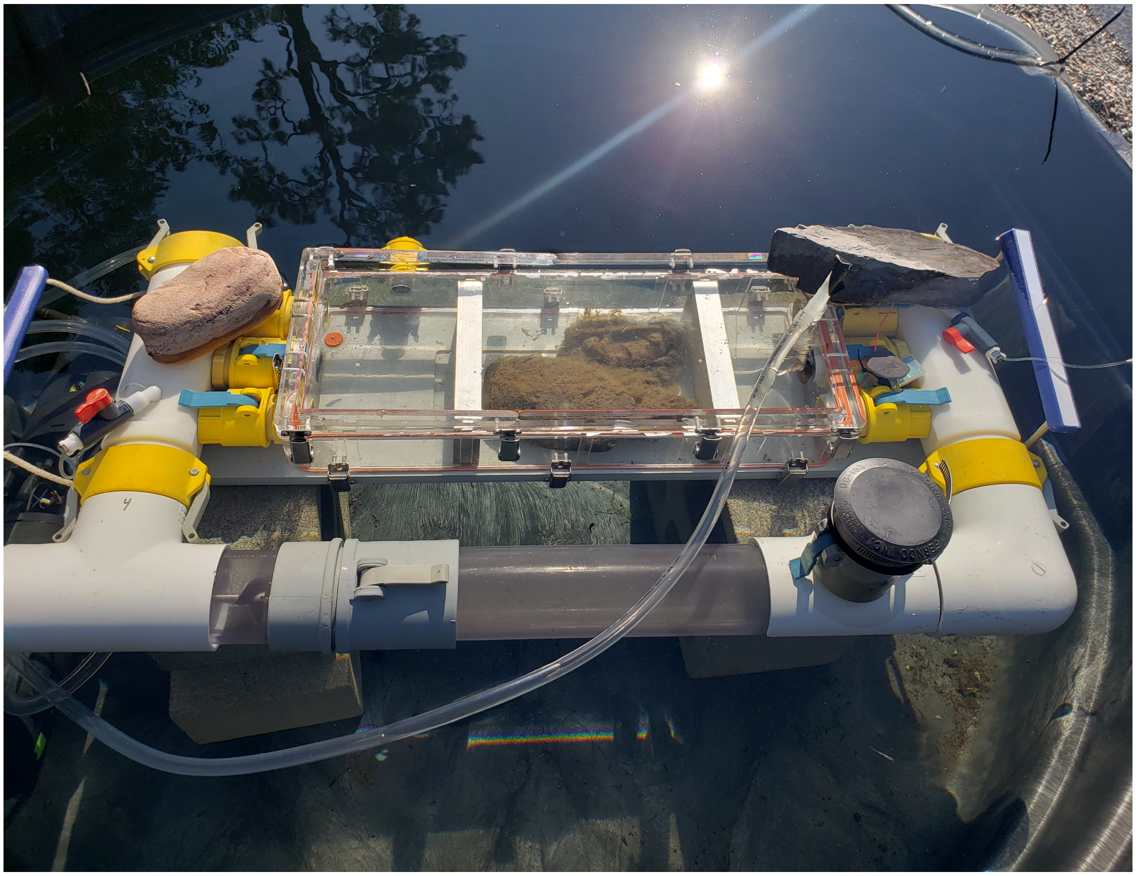

SI Figure 2: Example picture of a sealed and filled chamber used in this study. The chamber is submerged in lake water to limit temperature fluctuations. Water circulates through the white PVC arms from left to right. The clear tube emerging from the right of the chamber box is the return flow from the in-line dissolved gas sample system. The water for sampling was removed with the tube on the underside of the bottom left corner via a submersible pump. Not shown are the two sensors.
